# Plant electrophysiological reactions to specific environmental conditions

**DOI:** 10.1101/2024.12.03.626606

**Authors:** Soren H.P. Wilkening

## Abstract

Plant cells exhibit as much electrical activity as animal cells but the nature of these electrical variations remains poorly understood, particularly at the organism level. Transitions from darkness to light, mechanical shocks, temperature shocks, pathogen attacks, and physical damage to plants have been shown to elicit changes in plant electrical patterns at the organism level under laboratory conditions, however plant electrical behavior in natural environments remains comparatively understudied. Here, we demonstrate that simple extracellular voltage measurements across various wild plant species can identify associations between specific environmental conditions and patterns of electrical variation, ranging from seconds to hours, in plants exposed to these conditions.This work has potential applications in utlizing plants as environmental sensors.

## Introduction

Electrical signals in plants have been under investigation since their discovery in the 19th century. The study of plant electrophysiology has advanced in recent decades due to the application of biomolecular and machine learning techniques. Research has shifted from a focus on establishing the conditions for and the consequences of a specific plant electrical signal type that closely resembles the spiking of animal neurons [Trebacz 1998; Koselski 2008; Opritov 2005; Pikulenko 2005; Sukhov 2009/2011] and has taken a broader view by aiming to understand the full variety of fast and slow-onset plant electrical signal patterns [Mudrilov 2021; de Toledo 2019] in the context of other known signalling mechanisms [Szechyńska-Hebda 2017; Huber 2019; Yudina 2023], such as molecular transport, volatile organic compound diffusion, mechanical substrate wave [Mancuso 1999; Tran 2019] and acoustic wave propagation [Khait 2018] as well as electrical plant-to-plant signalling [Szechyńska-Hebda 2022]. In this study we aim to answer the question if plant internal state changes in reaction to specific environmental conditions can be identified and characterized by extracellular voltage measurements on plant stems.

## Materials and Methods

Six plant species and one control were recorded over periods ranging from 18 to 36 weeks (Table M1), three angiosperm species (Birch, Olive, Magnolia) and three gymnosperm species (Spruce, Larch, Yew) were chosen for this study. In addition environmental temperature, relative humidity and air pressure were recorded at each plant’s location.

### 1 Plant species

Spruce (Picea abies), wild, 10-15 years old, gymnosperm Larch (Larix decidua), wild, 5-10 years old, gymnosperm

Yew (Taxus baccata), potted outdoor, 3-5 years old, gymnosperm Birch (Betula pendula), wild, 5-10 years old, angiosperm

Olive (Olea europea), potted outdoor, 5-10 years old, angiosperm

Magnolia (Magnolia soulangeana), potted outdoor, 2-5 years old, angiosperm

### 2 Location and climatic conditions

All plants were located in the austrian Alps at altitudes between 800 and 1000 meters, within an area of approximately one square Kilometer. The wild plants at 1000 m altitude on a sunny, southward-facing side of a valley in the Lungau region, the potted plants at an altitude of 800 m. A brief description of the climatic conditions (values are averages): Hottest month July (18 C), coldest month January (-1 C), wettest month September (137mm), driest month January (39mm), annual precipitation 1012mm, average relative humidity 76%, average pressure 1017 mbar. See also Figure 1 and Figure 2.

**Figure 1.**
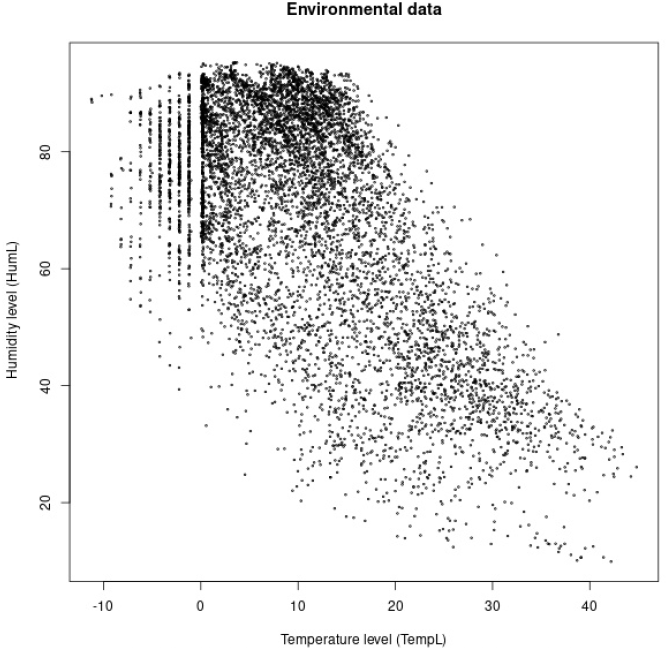
Temperature (C) and humidity (%) as measured at field location

**Figure 2.**
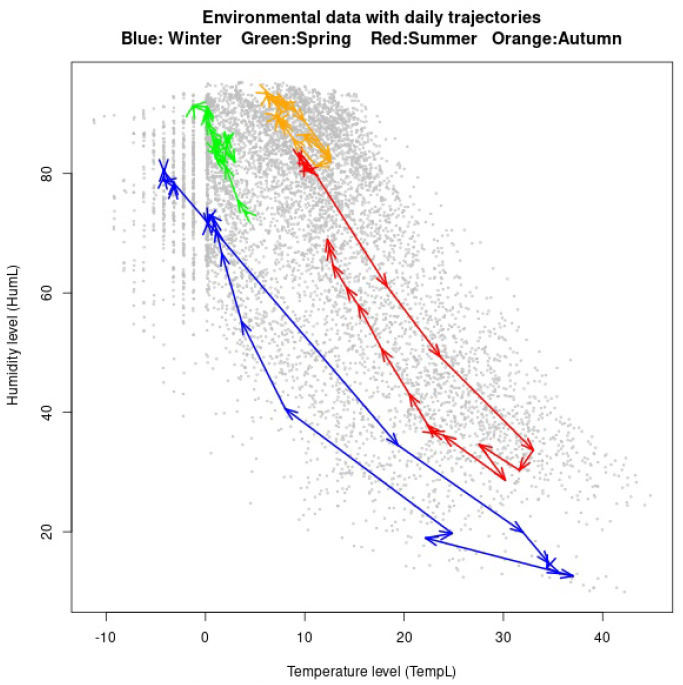
Temperature (C). humidity (%). Arrows indicate daily trajectories over time

### 3 Voltage measurement equipment and recording setup

The electrophysiological recording equipment consisted of one stainless steel needle electrode (E1) horizontally inserted into each plant’s stem at the height of the canopy, between approximately 40 cm and 2 m above ground depending on plant size, a second identical electrode (E2) inserted into a branch at the same height as the first electrode as well as a third reference electrode (E0) either inserted into the lower portion of the plant stem just above the ground or directly into the ground substrate. The two highest placed electrodes on each plant (E1,E2) were connected to the inputs of a high impedance, two-channel DC voltage amplifier (gain: 2-3) while the third electrode (E0) was connected to the amplifier’s reference input. This way the voltage difference between each stem/branch electrode and the reference electrode was measured for every plant continuously over a period ranging from 18 to 36 weeks depending on the plant (Table M1). In addition a control system was created and measured for a period of 10 consecutive weeks. It consisted of a dead branch 2 cm in diameter and 50 cm in length that had been cut from a Pavlovnia imperialis tree and left on a heap of wood cuttings for two years to ensure that no significant amount of live plant cells would be present at the time of the experiment. Immediately before the start of the control recordings this dead branch was inserted into a pot with gardening soil and connected to the electrodes E1 and E0.

### 4 Data acquisition and pre-processing

#### 4.1 Environmental data

Every plant/control recording location was equipped with a battery-powered environmental recorder separate from the battery-powered electrophysiology recorder to avoid electrical crosstalk. Using a sensor (BME680 Siemens) connected to a microcontroller (STM32) this device measured air temperature (Celsius), relative humidity (%) and air pressure (mbar) and stored the respective values on a microSD card once every 31 seconds.

#### 4.2 Plant and control electrophysiology data

On all plants and on the control system voltages were recorded from electrodes E1 and E2 at a sampling rate of 400 samples per second (sps). Timestamped voltage values in the range between -500 mV (millivolt) and +1000 mV were written to a microSD card by each recording device. Individual recording sessions lasted between 4 and 7 days with gaps of no more than 30 minutes between sessions in most cases. All recording devices were battery powered to enable field operation independent of the power grid and to reduce electrical noise to a minimum. To facilitate analysis all recorded voltage data was subsequently resampled to a final rate of 0.25 sps corresponding to one sample every four seconds.

### 5 Data processing

#### 5.1 Processing environmental data

The raw environmental data timeseries for each plant/control had three dimensions (Temperature, Humidity, Pressure) and one datapoint every 31 seconds. The 20%, 40%, 60%, 80% quantiles for each dimension were calculated using the entire length of the timeseries and were used for discretizing each dimension into five equally sized bins labeled ‘verylow’, ‘low’, ‘mid’, ‘midhi’ and ‘hi’ for humidity and similarly for the other two dimensions. (Figure 3)

**Figure 3.**
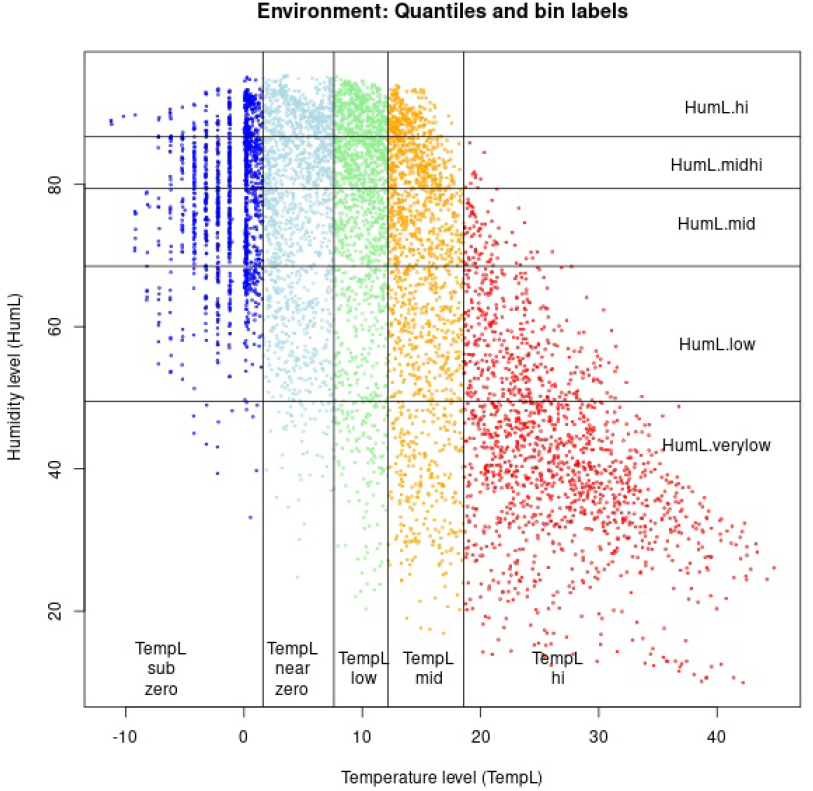

Each dimension was then aggregated into hourly intervals. The mean of all Temperature/ Humidity/ Pressure values within a given hourly interval was calculated and substituted by its respective bin label. In addition, the difference between the first and the last temperature/humidity/pressure value within a given hourly interval was calculated and substituted by its respective bin label, where the bins were calculated from the distribution of hourly differences (Figure 4 and Figure 5). These bins labeled ‘down2’, ‘down1’, ‘no’, ‘up1’, ‘up2’ indicated the speed of change for each of the three dimensions temperature/ humidity/ pressure in a given hour.

**Figure 4.**
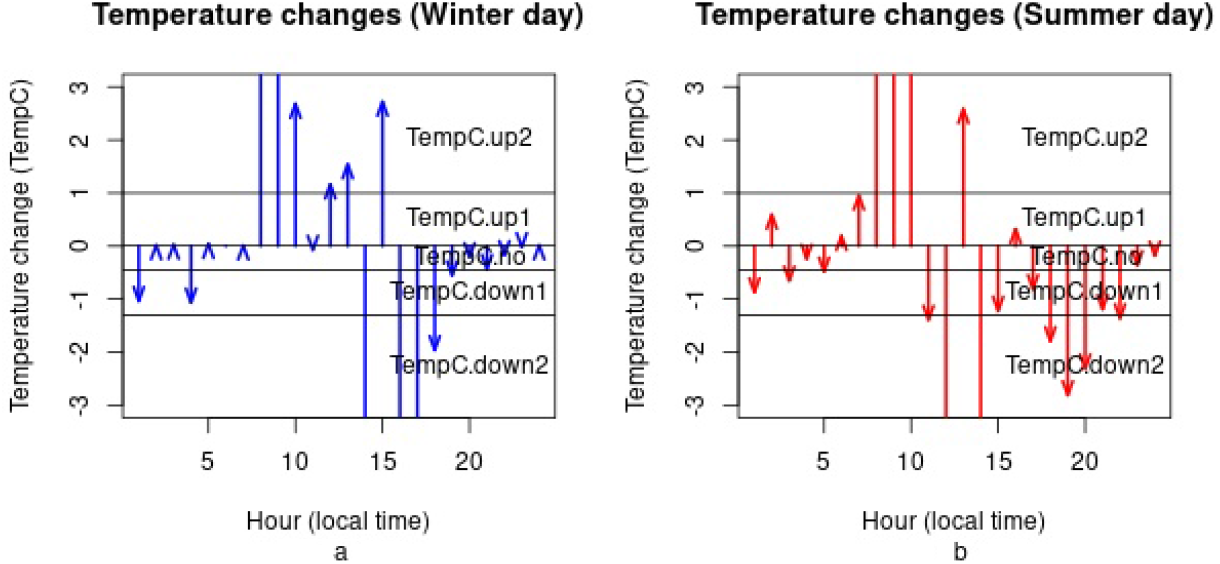

**Figure 5.**
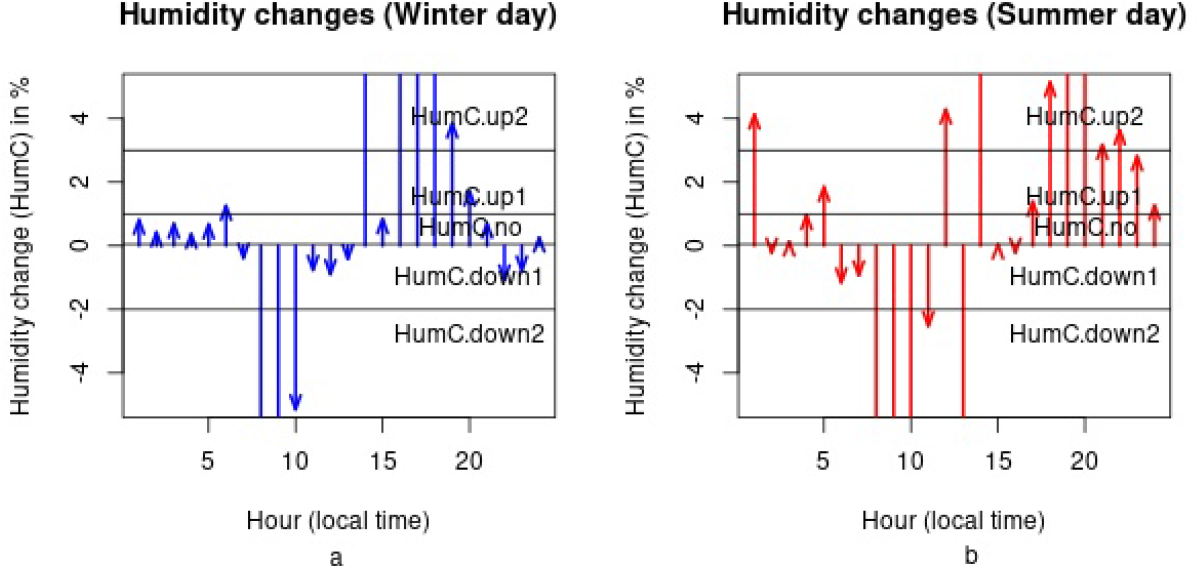

This discretization process produced an hourly environmental data timeseries of six dimensions that only contained the labels defined above. The final dimension names were defined as: Temperature level (TempL), temperature change (TempC), humidity level (HumL), humidity change (HumC), pressure level (PresL), pressure change (PresC).

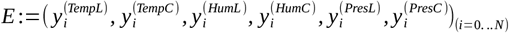

#### 5.2 Processing plant and control electrophysiology data

For a given plant/control the E1 electrode voltage timeseries *X* :=*xi*=0… *N* spanning the entire recording period was processed as follows. A constant c>0 was added to X to make all values positive. The resulting series was then normalized by componentwise logarithm and subtraction of the first component’s log to:

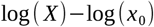

Next, a stepwise discrete fourier-transfom (DFT) without overlaps was calculated on the normalized X using a window length of 512 samples (∼ 34 minutes of data). Windows extending over the border of a recording session were discarded. For each window the DFT result vector of 512 complex values was first transformed to real values by using the complex Modulus, then the second half as well as the first value of each result vector were discarded, leaving a vector of length 255. Furthermore the logarithm was applied again to every vector component and by combining the output vectors of all windows, the data processing resulted in a multivariate timeseries of 255 dimensions with hourly timestamps for each plant/control where each dimension corresponded to a frequency bin of bandwidth 0.0005 Herz (Hz) in the frequency range between 0.5 milliherz (mHz) and 125 mHz. (Figure 8a-c, Figure 9a-c, Figure 10a-c)

**Figure 6.**
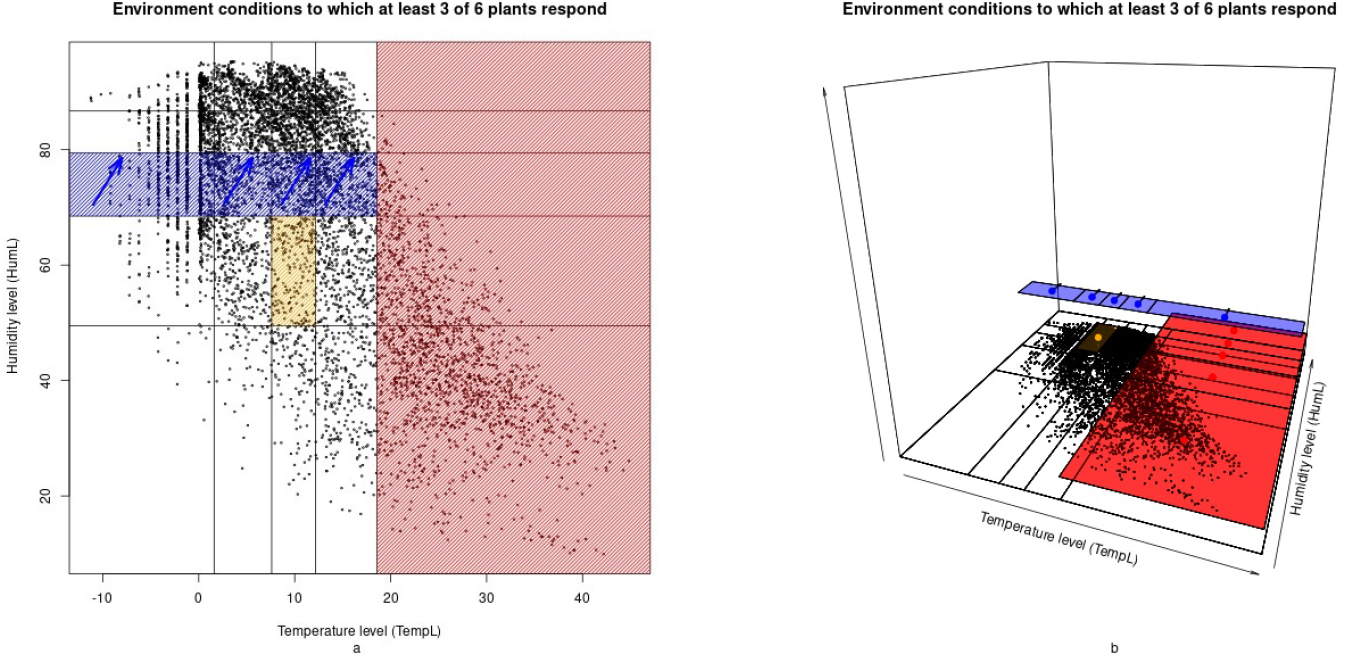

**Figure 7.**
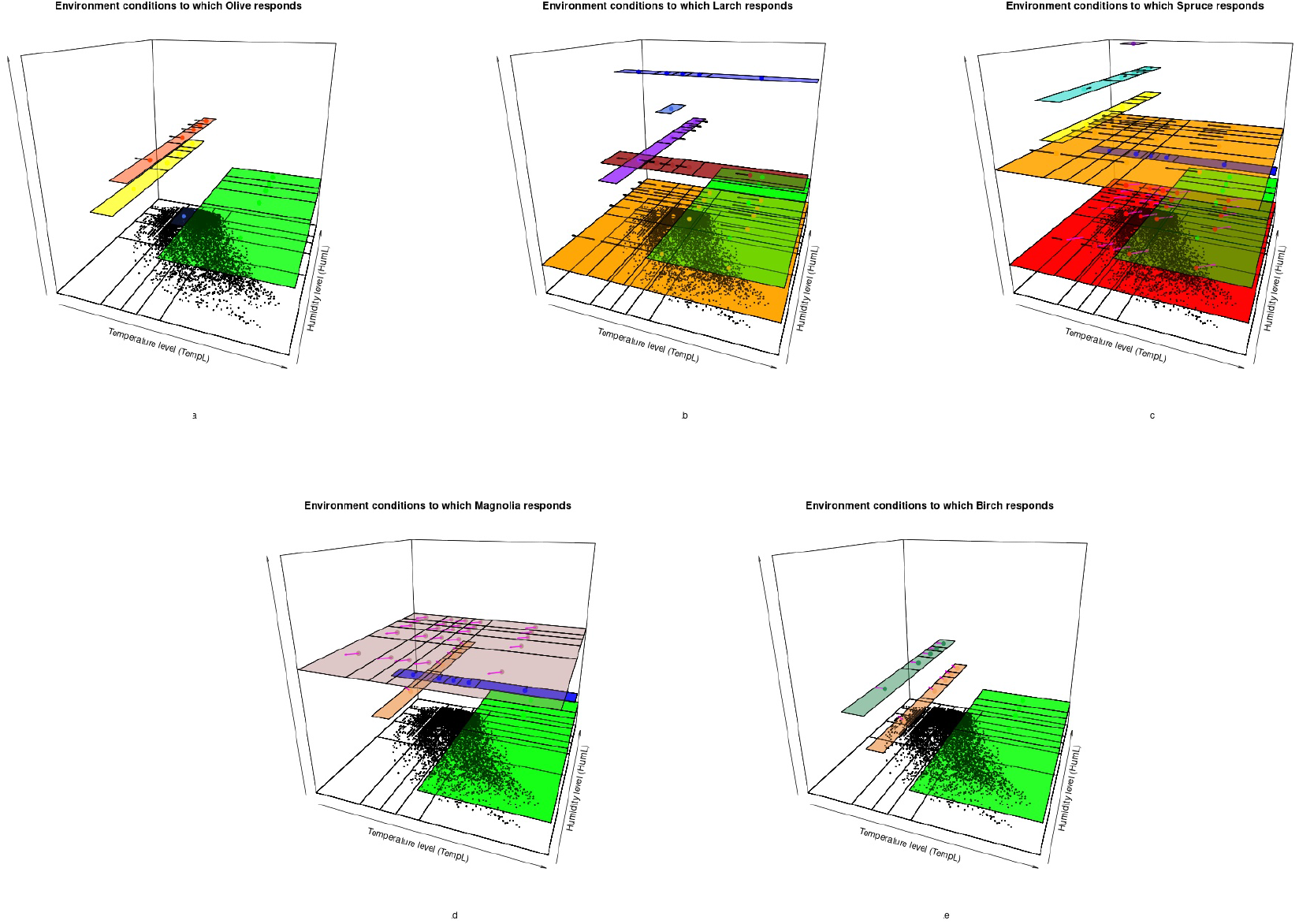

**Figure 8.**
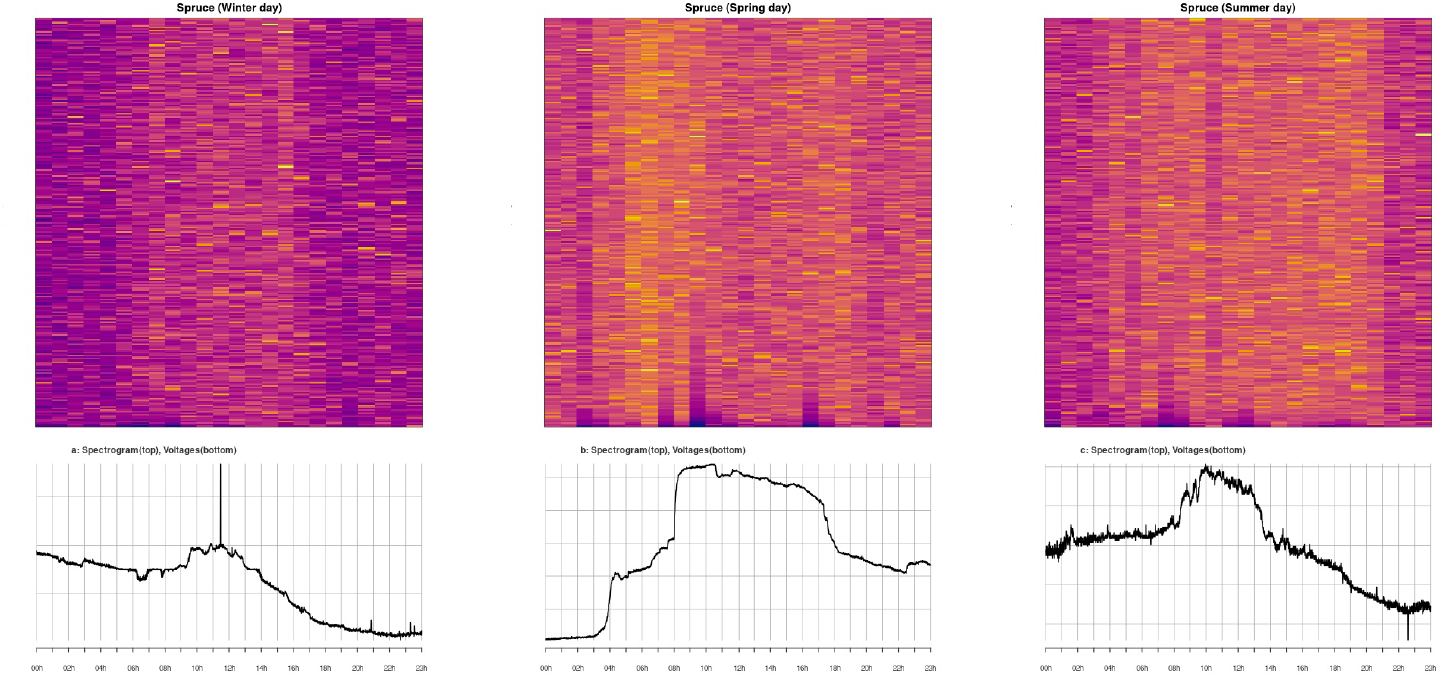

**Figure 9.**
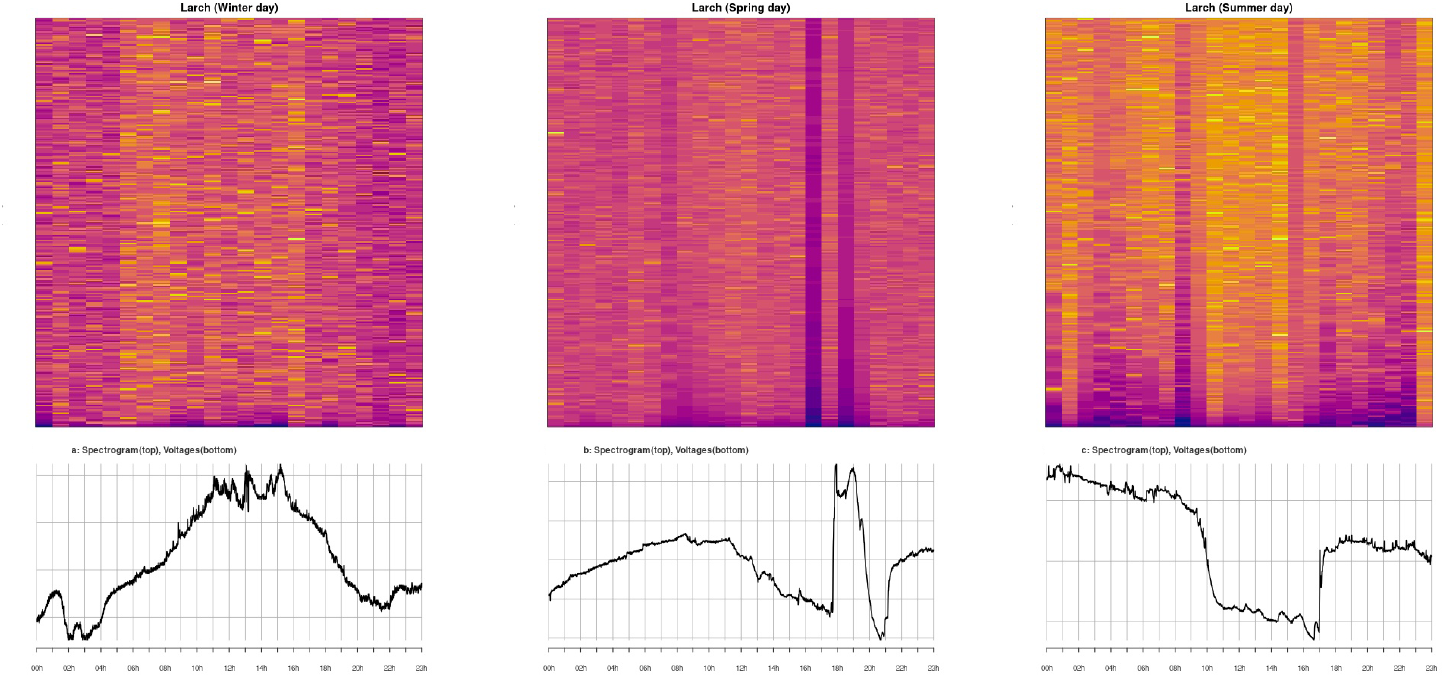

**Figure 10.**
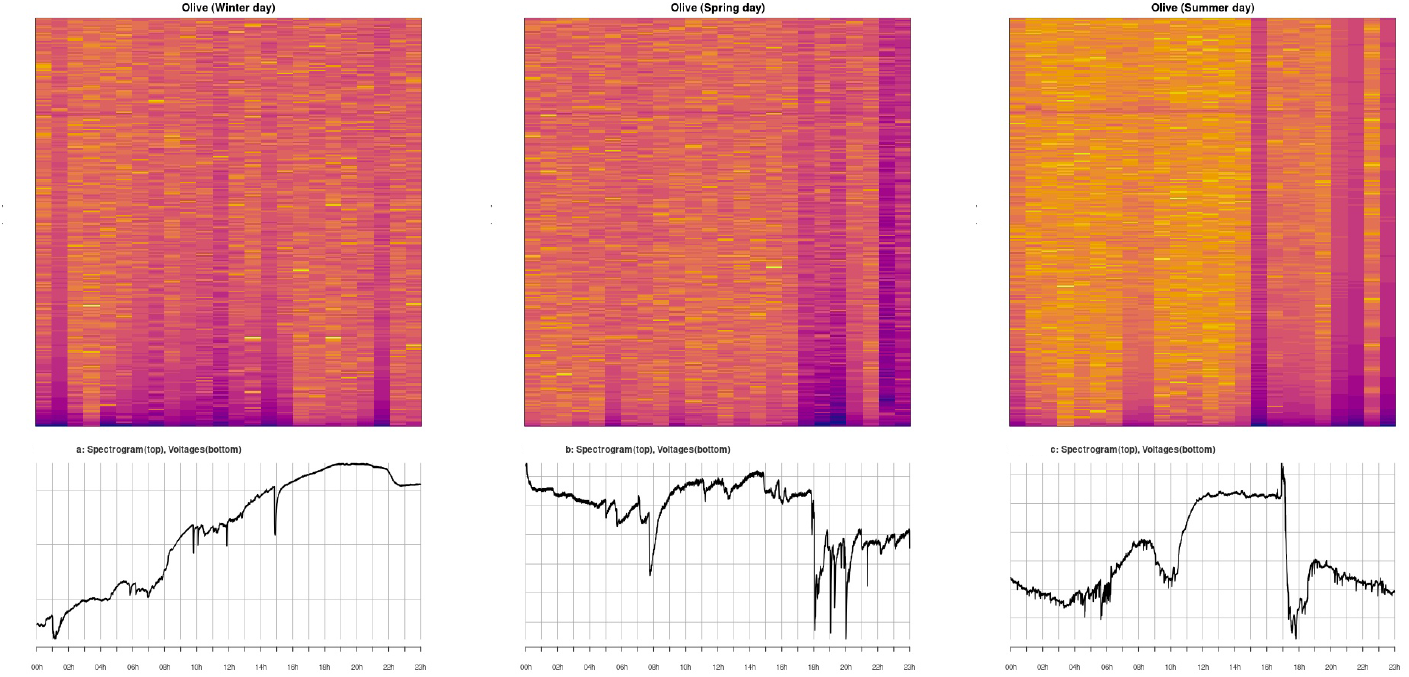

Blind source separation was then performed on the multivariate timeseries for every plant/control by applying Independent Component Analysis (R Version 4.2.2 statistical software; Package fastICA Version 1.2-5.1). The ICA method is predominantly used in neuroscience to separate multichannel electrical recordings into a smaller number of independent and maximally non-gaussian components for the purpose of dimensionality reduction. To determine a suitable number of target components heuristics were used to assess the featurespaces resulting from ICA decompositions for several values between 3 and 127 and the number of target components to use was fixed to 17. This resulted in a dimensionality-reduced representation of every plant/control by an hourly timeseries of 17 dimensions.

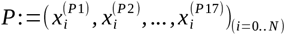

## 6 Analysis

Finding associations between environmental and plant electrophysiology data

For a given plant, the previous dataprocessing steps led to a multidimensional timeseries that characterized a plant’s state on an hourly basis through a numerical vector of length 17 (5.2). Likewise the environmental conditions the plant was finding itself in during the recording period were characterized by a six-dimensional timeseries of labels (5.1). To find associations between environment and plant/control we proceeded with fitting generalized linear models (R function glm) to the timeseries data.

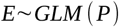

We used all the dimensions of the plant/control timeseries as independent variables and chose the dependent variable from the dimensions of the environmental timeseries. This approach may seem counter-intuitive, since we aim to predict environmental variables with plant variables. However, because of the richness of the plant latent space in comparison to the environmental latent space our chances of discovering meaningful associations between the two is greater if the analysis is conducted this way. If a strong association between a plant electrical pattern and a specific environmental condition existed it would be possible to find a well-fitting model with better-than-random predictive performance.

### 6.1 Motivation for using forward-looking analysis

When compared to animals, most plants’ reactions to environmental changes are slow and can take minutes or hours to manifest. In order to increase the chances of discovering delayed or slow-onset plant electrical reactions, our analysis of plant data took a forward-looking approach. When searching for assocations with a given environmental condition prevalent at time *t* _0_ we looked at plant data at time *t* _0_ but also included plant data for times *t* _0_ < *t*_1_ <*t* _2_ < *t*_3_ <*t* _4_ . Under the assumption that a plant may react to environmental change with an unknown delay, we would then be able to identify an association, if one existed.

### 6.2 Use of symbolic variable-value terms for describing subsets of environmental conditions

After binning (5.1) each of the six environmental dimensions had a range of five possible values. We chose to use simple (size 1) and complex (size 2) variable-value pairs to further subset the environmental conditions encountered during the experiment.

Examples of simple (size 1) variable-value terms (left) and their definition (right):

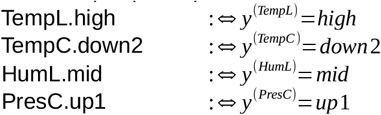

Examples of complex (size 2) variable-value terms and their definitions:

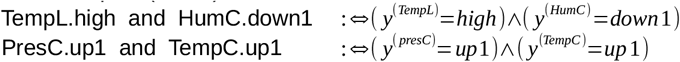

### 6.3 Fitting gl-models to discover associations between environmental and plant variables

Examples of possible associations described by simple left-hand-side (LHS) terms of size 1 are:

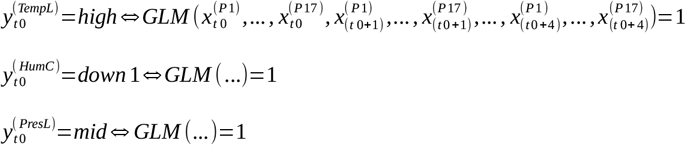

An example of an association described with a complex LHS term of size 2 is:

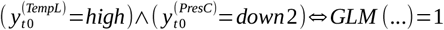

By our definition the dependent variable in these two types of terms was binary and consequently all fitted gl-models were binary classifiers. We tested all possible terms of size 1 (30 terms) and size 2 (375 terms) against all plants and control. Terms of size 3 or higher were not tested since the number of datapoints for each such term was too small to give significant results. For every term and plant/control combination we used out-of-sample k-fold validation (2<= k <= 5) to calculate a ROC/AUC value for the fitted gl-model. Briefly, the AUC (Area Under the Curve) measures the predictive performance of a binary classifier. AUC values fall between 0 and 1. A value of 0.50 or less indicates that the model performs no better than random and a value of 1.0 means that the models’ predictions are perfect. Values above 0.60 indicate acceptable, non-random predictive performance.

### 6.4 Selection of environmental subset terms most likely to reflect an association between plant and environment

If a specific environmental condition can be predicted by measuring voltage variations in a dead piece of wood exposed to these conditions, then finding that a live plant’s voltage variations can also predict the same enironmental condition is not a strong argument in favor of the plant voltages posessing unique predictive qualities when compared to an abiotic control system. On the other hand if an environmental condition cannot be predicted from a dead piece of wood (abiotic control system) but can be predicted from a live plant, this would support that the plant’s voltage variations do posess unique qualities associated with that environmental condition. We formalize this logic by defining three criteria that need to be met for any given (environmental condition) term in order to include it into the final selection of terms likely to be associated with plant electrical variation patterns. Criterion1: The AUC value for the term in the control experiment has to be less than 0.55 (Control predictive quality is random). Criterion2: The number of test cases in the control experiment has to be at least 50 (significance). Criterion3: The AUC for the plant experiments have to be greater than 0.65 (plant predictive quality is better than control predictive quality). Applying these criteria then led to the selection of the (environmental condition) terms in Table T1. Of special interest are those that have AUC values well above 0.65, have a small confidence interval radius and are present in the term selection for three or more of the six plants observed in this study. These terms have been separately listed in Table T2 and are visualized in Figure 6 (shaded areas).

## Results

An association between high temperature (>18 C) and a recognizable plant electrical pattern ocurring during the four hours following the onset of high temperatures was found (Figure 6, red shaded area). In addition an association between mid-level humidity (67-78 %) slowly increasing at a rate of 1.0-3.0 %/h and a recognizable plant electrical signalling pattern was found (Figure6, blue shaded area). Furthermore an association between low temperature (6-11 C) paired with low humidity(50-67 %) and a recognizable plant electrical signalling pattern was found (Figure6, orange shaded area).These associations were discovered in at least three different plants (Table T2).

For the six plants included in this study we found that our methodolygy was able to identify between three and eight high strength associations for each plant (Table T1) with the exception of the yew for which no associations were found. For spruce we identified eight associations, for larch six associations, for birch we identified three, for olive four and for magnolia also four associations (Figure 7a - 7e).

## Discussion

Our results establish a link between specific environmental conditions and time-delayed plant electrical patterns. We have used the expression ‘association’ to describe this link. We found three narrowly defined environmental conditions (Table T2) that elicit a characteristic electrical response in at least half of all the plants included in this study, suggesting that they might be detectable in the majority of angiosperm and gymnosperm species. Furthermore we have found environmental conditions that only some species repond to (Table T1).

Given that plants are known to react to and adapt to a variety of external stimuli a causal effect of the environment on the tested plants is the most likely explanation for the observed associations.

One of the strongest associations linked the onset of high temperature (> 18 C) to a subsequent electrical pattern. Research conducted by [Gilbert 2006] showed that the absolute diel plant electrical potential variation is strongly correlated to sapflux. Furthermore it is well established that sapflux is driven predominantly by the atmosphere-to-plant water vapor pressure deficit (VPD) [Wan 2023] which in turn is almost completely determined by air temperature and modulated by relative humidity [Grossiord 2020]. Therefore the association we found between high temperature and plant electrical variation is consistent with current research on sapflux and VPD dynamics and a purely thermoelectrical effect can be excluded due to the behavior of our control system which showed no association between high temperature and electrical pattern changes.

The yew was the only plant for which no significant association was found. Possible explanations are a fault in the plant data acquisition system or weak contacts between plant tissue and stem electrodes.

Our methodology used extracellular voltage measurements on plants in combination with data analysis techniques derived from neuroscience to non-destructively identify and characterize plant reactions and plant internal states. For future studies, substituting the general linear models with neural network models will likely improve the identification and characterization of environmental conditions and other external factors that impact a plant. We believe that our findings are a useful addition to the methods and tools available in plant biology in general and plant electrophysiology in particular since they enable researchers to use plant electrical reactions in near realtime for the characterization of otherwise difficult to observe plant state changes in response to external stimuli.

## Author contributions

S.W. conceptualized and designed the study, performed the field experiments, conducted the data analysis, interpreted the data, drafted and finalized the manuscript.

## Acknowledgements

S.W. wishes to thank the manager and the members of the Earth Species Project forum for fruitful discussions.

## Tables

**Table M1.** Plant species and recording periods.

| Plant | Duration of Recording | Date Start/End |
| --- | --- | --- |
| *Spruce (P. Abies)* | 32 weeks | Feb 2023/Oct 2023 |
| *Larch (L. Decidua)* | 19 weeks | Jan 2024/Jun 2024 |
| *Yew (T. Baccata)* | 36 weeks | Nov 2022/Aug 2023 |
| *Birch (B. Pendula)* | 20 weeks | Jan 2024/Jun 2024 |
| *Olive (O. Europea)* | 20 weeks | Mar 2024/Aug 2024 |
| *Magnolia (M. Soulangiana)* | 18 weeks | Apr 2024/Aug 2024 |
| CONTROL | 10 weeks | Sep 2024/Nov 2024 |

**Table T1.**
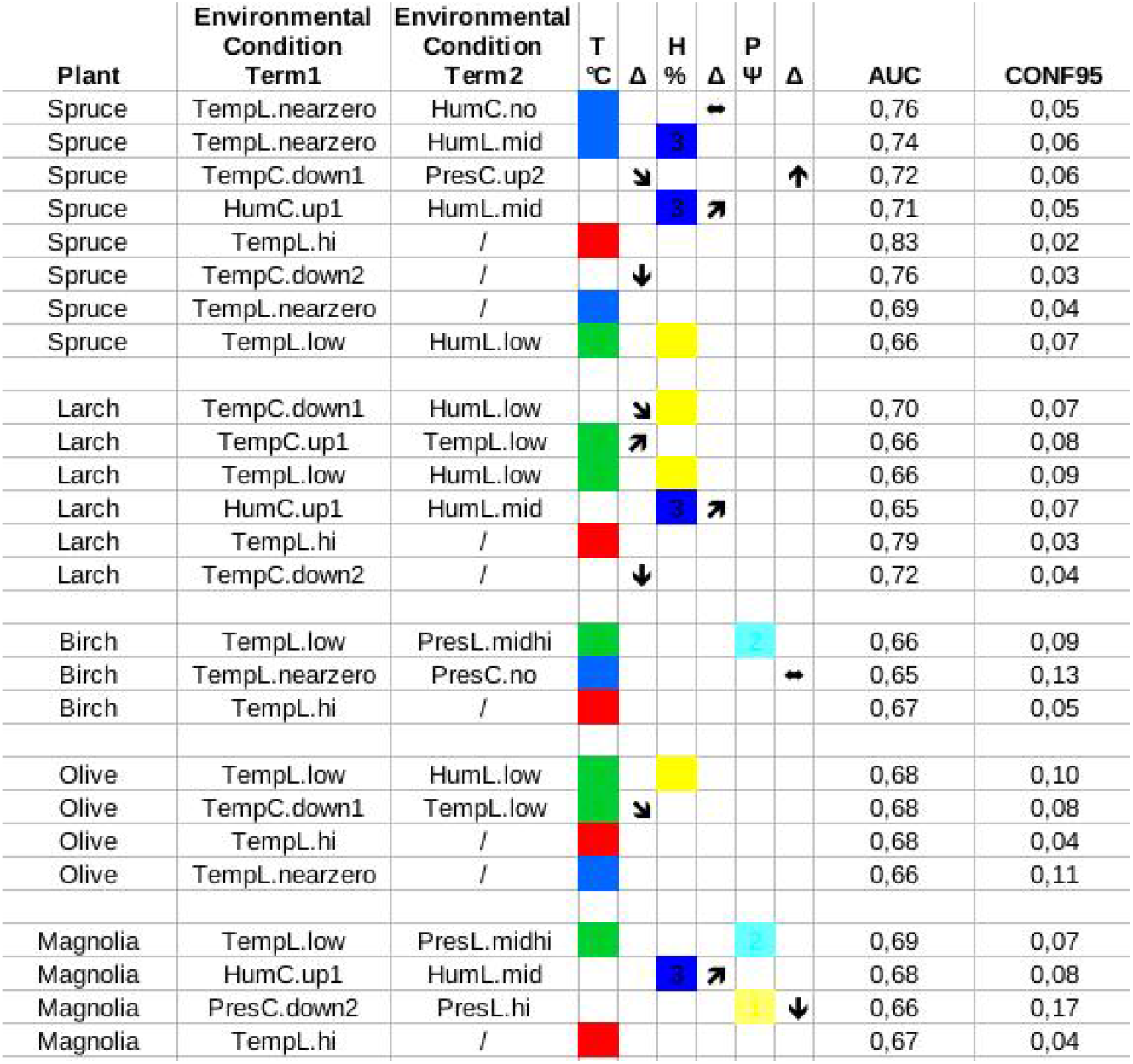
Environmental conditions to which at least one plant reacted.

**Table T2.**
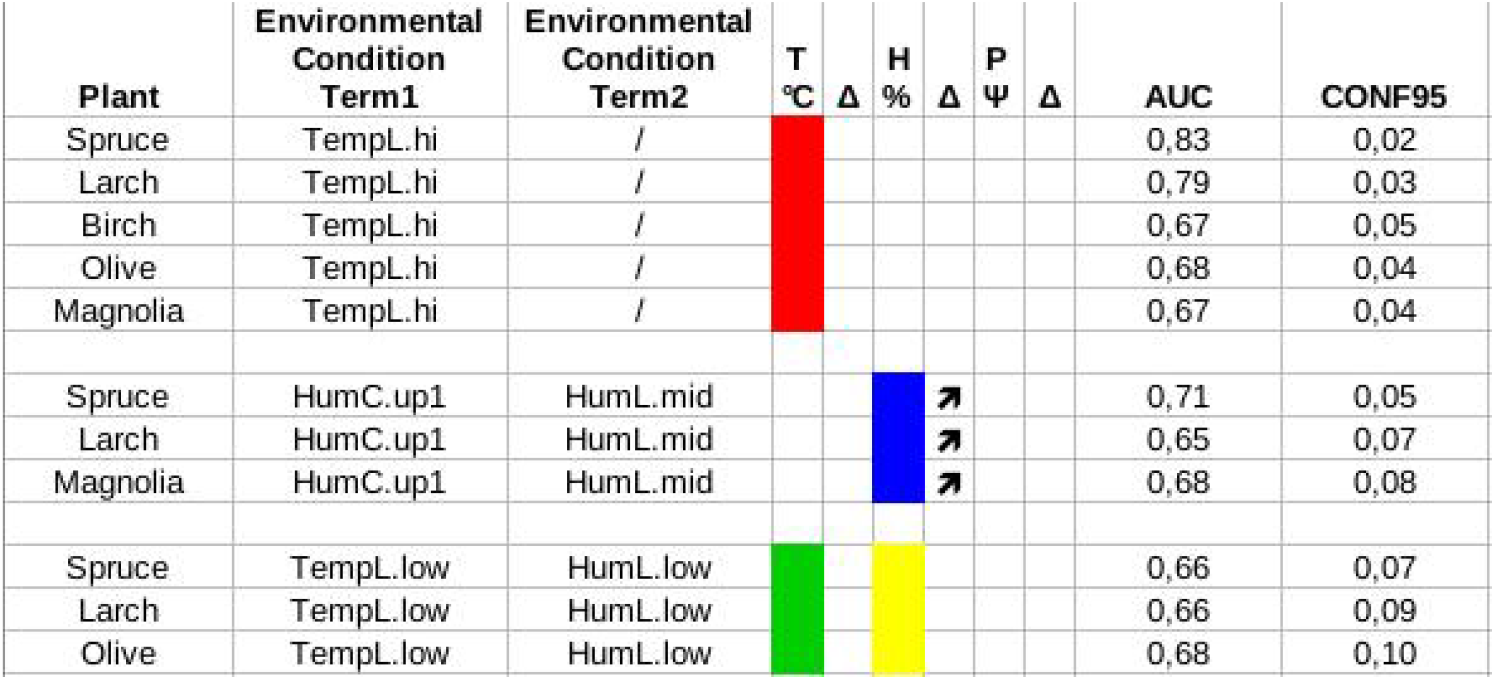
Environmental conditions to which at least three plants reacted.

